# MIRATS framework: Normative multiscale characterization of brain regulatory systems across sex and age using multimodal MRI

**DOI:** 10.64898/2026.06.15.732296

**Authors:** Zewu Ge, Weibei Dou

**Affiliations:** Department of Electronic Engineering, Beijing National Research Center for Information Science and Technology (BNRist), Tsinghua University, Beijing 100084, China

**Keywords:** multimodal MRI, brainstem, diencephalon, sex differences, aging, topological data analysis

## Abstract

Deep brain systems involved in arousal, autonomic regulation, sensory integration, and homeostatic control remain underrepresented in conventional whole-brain neuroimaging frameworks. In particular, diencephalic and brainstem nuclei are often insufficiently represented in cortex-centered analyses, limiting the normative references needed to interpret systems-level variation in health and disease.

To address this gap, we developed a unified multiscale framework with explicit representation of deep nuclei. By integrating cerebral, cerebellar, diencephalic, and brainstem atlases in standard space, we constructed a 220-region whole-brain parcellation and extracted complementary features at three analytical scales: nodal properties, edge-wise connectivity, and persistent-homology-based topological descriptors. We applied this framework to healthy adults from the Human Connectome Project-Aging cohort to characterize normative multiscale organization and test sex- and age-related variation.

Applied to this cohort, our framework revealed pronounced heterogeneity across anatomical systems. Brainstem and diencephalic nuclei showed multiscale feature profiles distinct from those of cerebral and cerebellar regions across nodal, edge-wise, and higher-order topological scales. Sex comparisons identified selective differences across different scales, whereas age modeling revealed widespread but feature- and system-dependent variation across adulthood. Together, these findings show that normative whole-brain organization in this deep-system-aware space is structured by system-specific rather than globally uniform patterns.

These findings establish a normative multiscale framework for characterizing brainstem-diencephalic-cerebellar-cerebral organization in healthy adults and provide a quantitative reference for future translational studies of disease-related abnormalities in deep regulatory systems.

## 1. Introduction

Adaptive behavior and physiological stability depend on continuous coordination between large-scale brain systems and subcortical and brainstem circuits that monitor internal bodily states and regulate arousal, autonomic function, sensory integration, and homeostatic balance [1, 2]. Within this distributed architecture, the diencephalon, brainstem, cerebellum, and cerebral cortex interact to support breathing, cardiovascular regulation, sleep-wake transitions, metabolic control, and the coupling of visceral states with cognition and behavior [3, 4, 5, 6]. Despite their central physiological importance, these deep brain structures and their distributed interactions remain incompletely represented in conventional multimodal neuroimaging frameworks, which have historically emphasized cortical organization and higher-order cognitive systems [7, 8, 9]. As a result, many whole-brain analyses provide only a partial account of the neural architecture underlying internal-state regulation.

Among these deep structures, the brainstem and diencephalon occupy especially important but methodologically challenging positions [10, 11]. The brainstem contains nuclei and circuits essential for respiratory rhythm generation, chemosensory processing, autonomic control, arousal modulation, and reflexive homeostatic responses [3, 10]. The diencephalon, including thalamic, hypothalamic, and related nuclei, contributes critically to sensory relay, state regulation, sleep-wake control, endocrine-autonomic coordination, and the integration of ascending and descending signals across the brain [1, 5, 6]. These structures maintain extensive reciprocal interactions with the cerebellum, subcortical nuclei, and cerebral cortex, forming a distributed network organization that cannot be adequately understood from a cortex-centered perspective alone [2, 4, 11]. However, due to the small size of many nuclei, limited contrast in standard-resolution magnetic resonance imaging (MRI), and the predominance of conventional cortical parcellation schemes, diencephalic and brainstem structures are frequently underrepresented or omitted in whole-brain analyses [8, 9, 12].

A second limitation concerns the analytical scale at which brain organization is described. Neuroimaging studies often examine morphometric, diffusion-derived, or functional measures separately [13, 14], or focus primarily on either regional features or pairwise connectivity [15, 16, 17, 18]. Yet brain organization is inherently multiscale. Local tissue properties characterize individual regions, interregional connections describe direct interactions, and higher-order topological structure captures global organizational principles that are not reducible to individual nodes or edges [19, 20]. Restricting analysis to only one of these levels may obscure important aspects of distributed organization, especially in systems in which deep nuclei, cerebellar territories, and cortical regions are tightly interdependent.

Topological data analysis (TDA), particularly persistent-homology-based approaches, offers a useful extension for addressing this problem [21, 22]. Unlike conventional graph measures that depend on a single threshold or emphasize only pairwise relationships, persistent homology characterizes the emergence and persistence of topological features across a filtration process [22]. In brain networks, such descriptors can quantify higher-order organizational properties that may not be fully captured by nodal or edge-wise features alone. Although TDA has been increasingly applied in connectomics [23, 24, 25], it is still rarely integrated as a formal component of a broader multiscale framework that also includes local tissue properties and interregional connectivity, especially in analyses with explicit anatomical representation of deep nuclei [21].

A third challenge is interpretive. Even when deep structures are included in whole-brain analyses, it often remains unclear how their imaging features should be interpreted in relation to normative brain organization. Diencephalic nuclei, brainstem nuclei, cerebellar regions, and cortical systems are unlikely to share uniform baselines in regional, connectional, or higher-order topological space. Without an explicit normative reference, observations made in disease may be difficult to contextualize. Establishing normative multiscale references is therefore not merely descriptive, but essential for biologically grounded interpretation of systems-level variation in brain organization [26, 27, 28].

These considerations motivate the need for a whole-brain framework with explicit representation of deep regulatory systems while also characterizing normative variability across key biological dimensions such as sex and age. Sex differences and age-related changes are among the most fundamental sources of inter-individual variation in brain structure and function, yet they have rarely been examined specifically in relation to deep nuclei within a unified multiscale framework. A framework capable of capturing such variation would provide not only a more complete account of healthy brain organization but also a stronger normative basis for future translational studies of disorders involving autonomic, respiratory, arousal, and homeostatic dysregulation [29, 30, 31, 32, 33].

In the present study, we developed a “Multimodal Integration of Regulatory Systems Across Topological Scales” (MIRATS) framework with explicit anatomical coverage of the cerebrum, cerebellum, diencephalon, and brainstem by integrating multiple standard-space atlases into a 220-region whole-brain parcellation. Within this common feature space, we derived complementary measures at three hierarchical scales: (i) nodal structural and microstructural properties; (ii) edge-wise connectivity features; and (iii) persistent-homology-based higher-order topological descriptors. We then applied the MIRATS framework to healthy adults to address three objectives: first, to characterize the normative multiscale organization of cerebral, cerebellar, diencephalic, and brainstem systems; second, to assess sex-related differences within this multiscale feature space; and third, to model age-related trajectories across adulthood. We hypothesized that deep nuclei would exhibit distinct normative profiles relative to other anatomical systems, and that the effects of sex and age would be expressed heterogeneously across analytical scales rather than uniformly across the whole brain.

## 2. Methods

Figure 1 illustrates the main modules and key concepts of the proposed MIRATS framework for characterizing normative whole-brain organization. The details of each component are described in the following sections.

**Figure 1:**
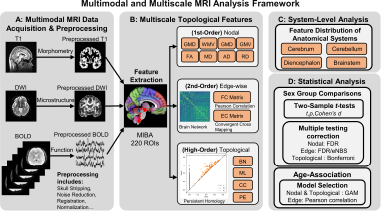
Overview of the proposed MIRATS framework. (A) Multimodal MRI data, including T1-weighted structural MRI, diffusion-weighted imaging (DWI), and resting-state blood-oxygen-level-dependent (BOLD) fMRI, were preprocessed to generate modality-specific measurements in a common analysis space. (B) Using the 220-region Multiscale Integrated Brain Atlas (MIBA), hierarchical features were extracted at three topological scales: first-order nodal features derived from morphometric and diffusion measures, second-order edge-wise features derived from functional connectivity (FC) and effective connectivity (EC), and higher-order topological features derived from persistent homology. (C) System-level normative analysis summarized multiscale feature distributions across the four major anatomical divisions: cerebrum, cerebellum, diencephalon, and brainstem. (D) Statistical analysis included sex-group comparisons with multiple-comparison correction and age-related trend modeling across feature scales. The MIRATS framework was designed to explicitly represent deep nuclei while preserving their large-scale interactions with cerebral and cerebellar systems. GMV, gray matter volume; WMV, white matter volume; GMD, gray matter density; WMD, white matter density; FA, fractional anisotropy; MD, mean diffusivity; AD, axial diffusivity; RD, radial diffusivity; FDR, false discovery rate; eNBS, extended network-based statistics; GAM, generalized additive model.

### 2.1. Dataset

Healthy participants were drawn from the Human Connectome Project-Aging (HCP-A) cohort, a large multimodal neuroimaging dataset designed to characterize healthy adult aging using harmonized MRI acquisition and phenotyping protocols [34]. Participants were included in the present study if they had available structural T1-weighted MRI, diffusion-weighted imaging (DWI), and resting-state blood-oxygen-level-dependent functional MRI (BOLD) of acceptable quality for multimodal feature extraction.

To address different analytical aims, we defined two partially overlapping healthy subsets. For sex-comparison analyses, a midlife subgroup of adults aged 36-55 years was used to reduce potential confounding from broader age heterogeneity while preserving sufficient sample size in both sexes. This subgroup included 320 participants (134 females and 186 males). For age-related analyses, a broader healthy adult cohort aged 36-80 years was used to model normative trends across adulthood. This cohort included 629 participants (354 females and 275 males). All analyses were based exclusively on de-identified public data acquired under HCP-A study procedures and approvals.

### 2.2. Multimodal MRI data and preprocessing

The acquisition protocol of multimodal MRI data is described in the original dataset documentation [35]. T1 and DWI data were obtained in raw form, whereas BOLD data had undergone minimal preprocessing pipelines [36] before our further preprocessing. Our preprocessing workflow was designed to generate modality-specific measurements in a common standard space suitable for multimodal joint analyses.

T1-weighted images were processed in SPM12 [37] running in MATLAB R2023b (MathWorks, Natick, MA, United States) to generate morphometric maps for regional tissue quantification. The preprocessing pipeline included skull stripping and tissue segmentation, resulting in gray matter and white matter probability maps.

DWI preprocessing was performed using the diffusion toolbox in FSL 6.0 [38] (FMRIB, Oxford, United Kingdom). The pipeline included brain extraction, eddy-current correction, motion correction, and tensor fitting. Voxel-wise diffusion-derived maps were generated from the fitted diffusion tensor model.

BOLD data were processed in DPABI V7.0 [39] running in MATLAB R2023b. Because minimal preprocessing had already completed head-motion correction, distortion correction, and spatial registration, the additional preprocessing in this study consisted of global signal regression and band-pass filtering (0.01-0.08 Hz). The resulting denoised time series were used for subsequent whole-brain functional analyses.

### 2.3. Whole-Brain Atlas Integration and Anatomical Organization

To enable unified whole-brain analysis with an explicit representation of deep nuclei, we constructed a whole-brain parcellation in standard space by integrating three atlases [40, 41, 42]. The resulting parcellation encompassed the cerebrum, cerebellum, diencephalon, and brainstem, thereby extending conventional cortex-centered templates to include deep structures relevant to arousal, autonomic regulation, sensory relay and integration, and homeostatic control.

For anatomical organization, all regions were assigned to four major divisions: cerebrum, cerebellum, diencephalon, and brainstem. Within the MIRATS framework, deep nuclei were operationally defined as the combined set of diencephalic and brainstem nuclei, including thalamic, hypothalamic, subthalamic, metathalamic, and related brainstem structures. This hierarchical organization enabled both fine-grained ROI-level feature extraction and higher-level system-based interpretation.

Ultimately, we generated a multiscale integrated brain atlas (MIBA) comprising 220 regions of interest (ROIs), including 98 cerebral regions, 26 cerebellar regions, 42 diencephalic regions, and 54 brainstem regions. The spatial coverage of the final template is presented in the Results section, and the abbreviations and descriptions of all 220 regions are provided in Supplementary Table 1. The MIBA resource is publicly available at https://github.com/gezw18/MIBA.

### 2.4. Multiscale Feature Extraction

Based on the MIBA-defined feature space, multimodal brain features were extracted at three hierarchical scales: first-order nodal features, second-order edge-wise features, and higher-order topological features. Our MIRATS framework was designed to capture complementary aspects of brain organization, ranging from local regional properties to pairwise interactions and network topology.

#### 2.4.1. First-Order Nodal Features

At the first-order scale, local morphometric and microstructural properties were quantified from the T1-weighted and DWI modalities. For morphometric features, the MIBA parcellation was transformed from standard space to each individual’s native T1 space to estimate regional gray matter volume (GMV) and white matter volume (WMV). In parallel, preprocessed T1-derived tissue maps were normalized to standard space, from which regional gray matter density (GMD) and white matter density (WMD) were calculated within the MIBA template. For diffusion microstructural features, all preprocessed diffusion-derived maps were normalized to standard space, and the mean values of fractional anisotropy (FA), mean diffusivity (MD), axial diffusivity (AD), and radial diffusivity (RD) were extracted for each MIBA region. In total, eight first-order nodal features were obtained.

#### 2.4.2. Second-Order Edge-Wise Features

At the second-order scale, interregional functional relationships were estimated from resting-state BOLD data to characterize pairwise coupling within the 220-region network. Specifically, the mean BOLD time series was extracted from each MIBA region. Undirected functional connectivity (FC) was computed as the Pearson correlation between regional time series, yielding a symmetric FC matrix that captured whole-brain temporal coupling. To characterize directed interactions, effective connectivity (EC) was estimated using convergent cross mapping [43], a nonlinear dynamical approach derived from state-space reconstruction theory that is suitable for inferring directional dependence beyond simple correlation structures. The resulting EC matrix provided a directed representation of interregional influence across the network. Together, FC and EC offered complementary descriptions of undirected coupling and directed interaction structures within the deep-nuclei-informed whole-brain network.

#### 2.4.3. Higher-Order Topological Features

To quantify network organization beyond nodal attributes and pairwise edges, persistent-homology-based TDA was applied to the whole-brain network and to anatomically defined subnetworks. Persistent homology tracks the birth and death of topological features across a range of filtration thresholds, thereby providing a threshold-free characterization of multiscale network organization [44].

From the resulting persistence patterns, we extracted four summary descriptors: Betti number (BN), reflecting the number of loops in the network; maximum lifetime (ML), reflecting the persistence of the most stable loop; persistent entropy (PE) [45], reflecting the complexity of loop organization; and the first Carlsson coordinate (CC) [46], representing a weighted persistence-based summary of the loop structure. The full Carlsson coordinate vector is four-dimensional, but only its first component was retained here as a compact topological fingerprint. These TDA features were computed separately for five networks, namely the whole-brain network and the four major anatomical subnetworks, under both undirected FC and directed EC representations.

### 2.5. System-Level Analysis of Normative Brain Organization

In addition to ROI-level feature extraction, our MIRATS framework was designed to support system-level characterization of normative brain organization. Specifically, regional features and connectivity patterns were summarized according to the major anatomical divisions defined above, allowing comparison of multiscale feature distributions across the cerebrum, cerebellum, diencephalon, and brainstem.

This system-level analysis served two purposes. First, it enabled evaluation of whether deep nuclei occupy distinctive positions in the healthy multimodal feature space relative to other anatomical systems. Second, it provided an interpretable normative reference against which future disease-related abnormalities may be contextualized. In this sense, the normative component of the MIRATS framework was not treated as an auxiliary analysis, but as an integral part of the methodological design.

### 2.6. Statistical Analysis

Because the primary aim of the present study was to establish a normative multiscale characterization framework in healthy adults, the analyses focused on two complementary sources of biological variation: sex differences and age-related trends. All statistical analyses were performed using Python-based scientific computing libraries.

#### 2.6.1. Sex Comparison Analysis

Sex-related differences were examined in the midlife subgroup (36-55 years) using independent-samples two-sided t tests. For first-order nodal features, statistical tests were performed for each ROI-feature combination. For second-order edge-wise features, tests were conducted for each FC and EC connection. For higher-order topological features, tests were conducted for each topological summary metric. To control false positives across the multiscale feature space, multiple-comparison correction was performed separately within each analytical scale. Specifically, the Benjamini-Hochberg false discovery rate (FDR) [47] procedure was applied to first-order nodal features and second-order edge-wise features. For edge-wise analyses, in addition to FDR correction, extended network-based statistics (eNBS) [48] were used to identify connected components showing sex-related effects. Because the number of higher-order topological measures was relatively small, Bonferroni correction was applied at this scale. A two-sided FDR-corrected *q <* 0.05 was considered statistically significant for first-order nodal and second-order edge-wise analyses, and a Bonferroni-corrected *p <* 0.05 was considered statistically significant for higher-order topological features.

#### 2.6.2. Age-Related Trend Analysis

Age-related trends were examined in healthy adults aged 36–80 years. Given the differences in feature dimensionality across scales, age effects were modeled using generalized additive models (GAM) for first-order nodal features and higher-order topological features, whereas Pearson correlation analyses were used for second-order edge-wise features. To minimize potential confounding by sex, all age-related analyses were performed separately in males and females.

For first-order nodal features and higher-order topological features, age trajectories were visualized using a two-stage GAM procedure. First, a GAM was fitted to the observed feature values as a smooth function of age to estimate the mean trajectory. Predicted mean curves and their pointwise 95% confidence intervals were then obtained on an evenly spaced age grid spanning the observed age range. Second, residuals from the fitted mean model were squared and modeled as a smooth function of age using a second GAM with the same spline specification. The fitted values from this residual-squared model were treated as smooth estimates of age-varying residual variance. Predicted variance values and their confidence bounds were constrained to be non-negative when necessary, and age-varying standard deviation trajectories were obtained by taking the square root of the fitted variance curve. Pointwise 95% confidence intervals for the standard deviation trajectories were derived by applying the same square-root transformation to the lower and upper confidence bounds of the variance model. These standard deviation trajectories should therefore be interpreted as model-based descriptive summaries of age-dependent dispersion.

Because these analyses were intended to characterize normative age-associated patterns rather than to establish a formally corrected set of age-sensitive features, no multiple-comparison correction was applied. Accordingly, age-related findings are reported as nominal or descriptive associations and should be interpreted cautiously.

## 3. Results

### 3.1. Normative multiscale organization of anatomical systems

#### 3.1.1. Structural and microstructural organization

We first assessed whether the four major anatomical divisions occupy distinct positions in normative structural and microstructural feature space, with particular attention to the deep anatomical systems. Eight regional metrics derived from T1-weighted and diffusion MRI, including gray matter volume, gray matter density, white matter volume, white matter density, fractional anisotropy, mean diffusivity, axial diffusivity, and radial diffusivity, were compared across the cerebrum, cerebellum, diencephalon, and brainstem (Figure 2). Across these measures, the kernel density distributions showed clear system-dependent stratification rather than a single shared profile.

**Figure 2:**
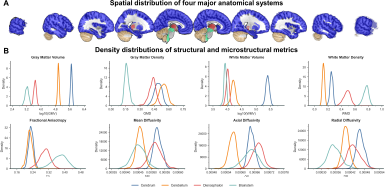
Schematic diagram of anatomical systems and density distributions of structural and microstructural metrics across subjects. (A) Spatial visualization of the four major anatomical systems, including the cerebrum, cerebellum, diencephalon, and brainstem. Each color denotes one anatomical system. (B) Kernel density distributions of regional metrics across the four anatomical systems.

Several robust trends were evident. The cerebrum occupied the highest GMV and WMV ranges, consistent with its dominant contribution to overall brain volume. The cerebellum, although smaller in volume, showed comparatively high gray matter density. In contrast, the brainstem displayed a particularly distinctive microstructural profile, with the highest white matter density and FA values, together with relatively low radial diffusivity. This pattern is consistent with the brainstem’s role as a major conduit for ascending and descending long-range white matter pathways. The diencephalon also showed a separable deep-system profile, generally occupying intermediate but shifted ranges across both morphometric and diffusion-derived features, including higher anisotropy-related values than the cerebrum and cerebellum. More broadly, diencephalic and brainstem regions tended to cluster more tightly and to occupy different parts of the feature space than cerebral and cerebellar regions, indicating that these deep systems have distinct structural and microstructural profiles. These findings suggest that deep anatomical systems are not simply part of a general non-cortical background, but instead represent differentiated structural components of whole-brain organization. Together, these results provide a normative anatomical framework for interpreting the system-specific functional and topological properties of deep brain nuclei.

#### 3.1.2. Functional and topological organization

We next examined whether this anatomical differentiation was also reflected in functional interactions and higher-order network architecture. Group-mean functional connectivity (FC) and effective connectivity (EC) matrices revealed clear system-dependent organization (Figure 3A). The FC matrix showed a pronounced block structure, indicating coherent within-system coupling together with structured between-system interactions. This pattern was also evident in the FC edge-weight distributions: the cerebellum was shifted toward stronger positive FC values, whereas the cerebrum showed the broadest distribution spanning both weak negative and positive correlations. In contrast, the diencephalon and the brainstem displayed narrower FC distributions concentrated around weak positive values, indicating more constrained functional coupling profiles in these deep systems. EC showed a similarly stratified but more uniformly positive pattern. Compared with the cerebrum, cerebellar EC weights were shifted toward stronger values, whereas the diencephalon and brainstem showed tighter distributions centered at relatively lower EC strengths. Thus, both FC and EC separated the four anatomical systems, with the diencephalon and brainstem again occupying distinctive positions within whole-brain communication architecture.

**Figure 3:**
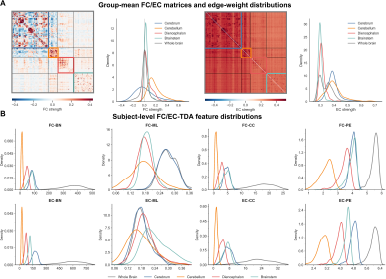
Group-level FC/EC networks and subject-level TDA feature distributions. (A) Group-mean FC and EC matrices and their edge-weight distributions. Colored boxes indicate anatomically defined within-system subnetworks. Density curves show the distribution of group-mean connection strengths for the whole brain and each anatomical system. (B) Subject-level distributions of FC-TDA and EC-TDA features across anatomical systems. Four TDA descriptors were examined: BN, Betti number; ML, maximum lifetime; PE, persistent entropy; and CC, Carlsson coordinate. Density curves represent inter-subject distributions of each feature for the whole brain and individual anatomical systems.

We then asked whether these system-level differences extended beyond pairwise connectivity to topological organization. Subject-level TDA features derived from both FC and EC showed clear distributional separation across the cerebrum, cerebellum, diencephalon, brainstem, and whole-brain network (Figure 3B). As expected, the whole-brain network occupied the highest ranges for several features, including BN, CC, and PE. Among the anatomical systems, the cerebrum generally showed relatively high BN values, whereas the cerebellum occupied lower ranges for BN, CC, and PE. The diencephalon and brainstem exhibited distinct deep-system signatures rather than simply following cerebral or cerebellar patterns: the diencephalon tended to occupy intermediate ranges across features, whereas the brainstem showed relatively elevated values for selected metrics such as FC-CC, FC-PE, and EC-ML. Collectively, these findings indicate that normative brain organization is structured across multiple scales, with the diencephalon and brainstem emerging as distinct components in both functional coupling and higher-order topology.

### 3.2. Sex-related multiscale differences in the midlife subgroup

We next examined sex-related variation in healthy midlife adults (36-55 years) within the normative multiscale framework. Sex effects were assessed across regional structural and microstructural features, pairwise connectivity, and TDA-derived topological measures, with Cohen’s d defined as male minus female.

#### 3.2.1. Sex-related differences in structural and microstructural features

Sex-related differences were observed across all four anatomical systems, but their extent and direction varied markedly by feature (Figure 4A; Supplementary Table 2). Among the eight measures, GMV and WMV showed the broadest and most consistent sex effects across the cerebrum, cerebellum, diencephalon, and brainstem, indicating widespread sex-related variation in regional morphology. Boxplots for GMV and GMD are shown in Figure 4B-C, and the corresponding boxplots for the other six features are provided in Supplementary Figure 1.

**Figure 4:**
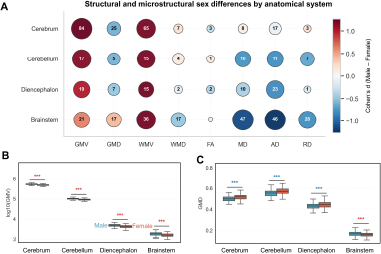
System-level summary of sex differences in structural and microstructural brain features. (A) Bubble plot summarizing significant sex differences across four anatomical systems and eight brain features. The number inside each bubble indicates the number of ROIs showing significant sex differences within the corresponding anatomical system and feature. Bubble size represents the proportion of significant ROIs relative to the total number of ROIs in that system. (B, C) Representative boxplots showing sex differences in two typical features: GMV and GMD (red, Male > Female; blue, Male < Female). \*\*\**q <* 0.001. The box bounds indicate the 25th and 75th percentiles and the center line indicates the median, whiskers correspond to minimum and maximum data values

A notable pattern emerged for gray matter metrics. In the cerebrum, cerebellum, and diencephalon, GMV and GMD showed opposite sex-related trends: GMV was generally higher in males, whereas GMD was generally higher in females. In contrast, the brainstem showed the same directional pattern for both GMV and GMD, with both measures tending to be higher in males. Thus, sex differences in gray matter organization were not uniform across anatomical systems, but instead followed a division-specific pattern that distinguished the brainstem from the other three systems. Diffusion-derived measures further highlighted the prominence of deep systems. FA showed relatively limited sex differences, whereas MD and AD exhibited more extensive effects, particularly in the diencephalon and brainstem. RD differences were more moderate overall but were again most evident in the brainstem. Together, these results indicate that sex-related variation in midlife adults is feature-specific and anatomically heterogeneous, with especially pronounced effects in deep anatomical systems rather than in the cerebrum alone.

#### 3.2.2. Sex-related differences in functional and topological organization

We next examined whether sex-related effects were also evident in functional connectivity and higher-order network topology. The statistical results at the edge-wise scale are shown in Figure 5A and Supplementary Table 3. Corrected sex differences in FC were confined to within-system connections of the cerebrum and cerebellum. These FC effects included both positive and negative differences, indicating that sex-related variation in undirected functional coupling was bidirectional rather than characterized by a uniform shift in one direction. In contrast, corrected sex differences in EC were distributed more broadly across the whole brain, involving connections spanning multiple anatomical systems, and all significant EC effects were positive (male > female), indicating a directionally consistent sex effect in effective connectivity. Thus, FC and EC showed clearly distinct sex-difference patterns, with FC effects restricted to intra-cerebral and intra-cerebellar organization, whereas EC effects were more spatially distributed and directionally consistent. We further used eNBS to re-examine significant connected components showing sex differences in FC and EC (Supplementary Figure 2). The resulting components involved connections distributed across the whole brain, indicating that sex-related differences were expressed at the level of distributed network organization rather than as isolated pairwise effects.

**Figure 5:**
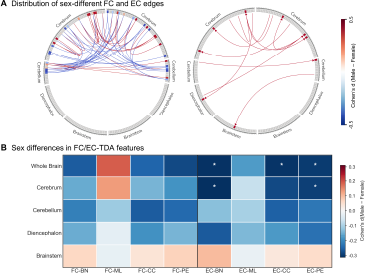
Sex differences in functional connectivity (FC), effective connectivity (EC), and TDA-derived features in the HCP-A cohort. (A) Distribution of FC and EC edges showing FDR-significant sex differences. The left and right circular plots represent FC and EC, respectively. (B) Heatmap of Cohen’s d values for sex differences in FC-TDA and EC-TDA features across the whole brain and the four anatomical systems. White asterisks indicate features that remained significant after Bonferroni correction (* Bonferroni-corrected *p <* 0.05).

At the higher-order scale, TDA-derived features also showed sex-related variation (Figure 5B; Supplementary Table 4). Across most systems and descriptors, effect sizes were predominantly negative, indicating that males generally showed lower TDA feature values than females. This tendency was especially evident for BN, CC, and PE, whereas ML showed weaker and less consistent differences across systems. Corrected effects were more prominent for EC-based than for FC-based TDA features, with the strongest and clearest differences concentrated in the whole-brain and cerebral networks. By comparison, cerebellar, diencephalic, and brainstem TDA differences were more selective. Even so, the effect-size patterns remained anatomically informative: the diencephalon tended to show negative shifts across several descriptors, whereas the brainstem showed modest positive shifts for several features. Boxplots for TDA features showing significant sex differences are provided in Supplementary Figure 3.

### 3.3. Age-related trends across adulthood

We next characterized age-related variation across the broader healthy adult HCP-A sample spanning adulthood (Figure 6A). Age trajectories were examined for whole-brain structural and microstructural measures, as well as for connectivity- and topology-related network features, with sex-stratified fits shown for descriptive comparison.

**Figure 6:**
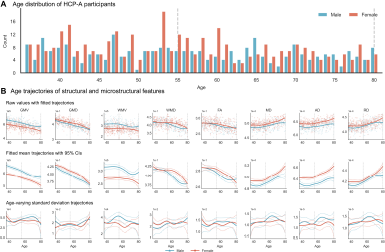
Age distribution and age trajectories of whole-brain structural and microstructural features in the HCP-A cohort. (A) Age distribution of HCP-A participants, shown separately for males and females. (B) Age trajectories of eight whole-brain structural and microstructural features. The top row shows individual raw values with fitted trajectories; the middle row shows fitted mean trajectories with pointwise 95% confidence intervals; and the bottom row shows age-varying standard deviation trajectories with pointwise 95% confidence intervals. Blue and red curves represent males and females, respectively.

#### 3.3.1. Age-related trends in structural and microstructural features

Whole-brain structural and microstructural features showed clear but feature-specific age trajectories (Figure 6B). GMV, GMD, WMD, and FA generally declined with age in both sexes, whereas MD, AD, and RD showed nonlinear age-related increases, characterized by relative midlife stability or slight early decreases followed by more pronounced upward shifts in later life. Compared with these measures, WMV showed relative stability across much of the observed age range and exhibited a modest late-life decline. Across most features, male and female trajectories followed similar age-related patterns but remained separated in overall level throughout adulthood, indicating stable sex-related differences alongside shared aging effects.

The corresponding trajectories for the four major anatomical systems are shown in Supplementary Figure 4. The cerebrum, cerebellum, and diencephalon broadly recapitulated the whole-brain pattern, with age-related decreases in GMV, GMD, WMD, and FA together with later-life increases in diffusivity measures. The brainstem, however, showed a partially distinct profile. Brainstem GMV still declined with age, and WMD also showed a downward trend, but GMD and FA were comparatively stable and more nonlinear than at the whole-brain level. In addition, age-related increases in brainstem MD, AD, and RD were attenuated and more sex-dependent than in the whole-brain results: these diffusivity measures increased more clearly in females, whereas male trajectories were comparatively flat and did not show the same late-life upward shift.

#### 3.3.2. Age-related trends in connectivity and topological organization

Age-related trends were also evident in FC, EC, and TDA-derived features. Edge-wise FC and EC analyses were performed separately in males and females, with FC shown in Figure 7A and EC in Supplementary Figure 5. In both sexes, age-associated edges involved not only intra-cerebral connections but also links between the cerebrum and the cerebellum, diencephalon, and brainstem, indicating broadly distributed age-related changes across cortical and deep brain systems. FC showed both positive and negative associations, with negative effects visually predominating, whereas EC effects were predominantly positive. Overall, males and females exhibited similar spatial distributions of age-associated connectivity patterns, with nominal associations in both groups involving comparable intra-cerebral and cerebrum-subcortical connections.

**Figure 7:**
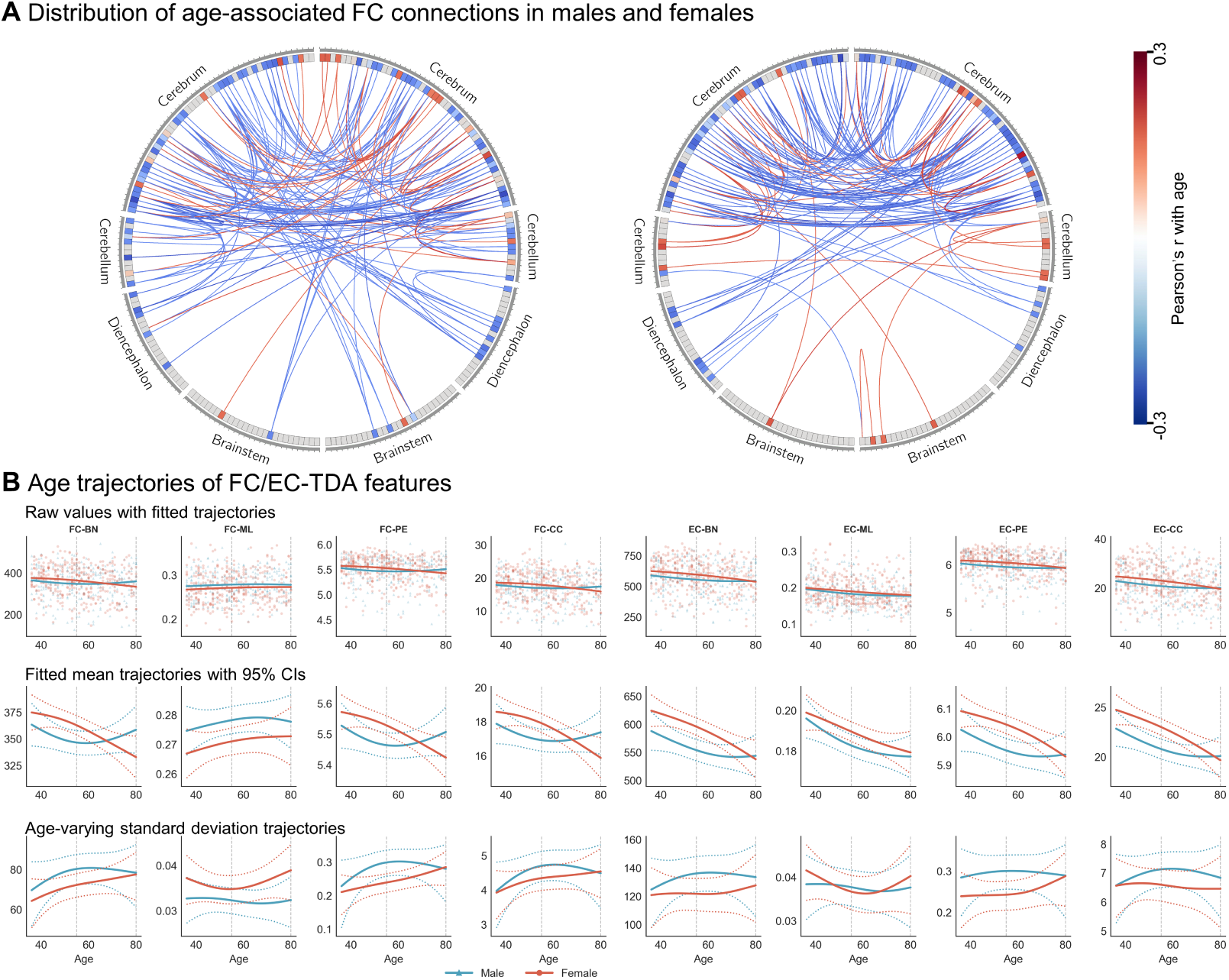
Age-associated connectivity patterns and TDA-derived age trajectories in the HCP-A cohort. (A) Spatial distribution of FC edges exceeding the display threshold of *|Pearson^′^s r| >* 0.2 , displayed separately for males (left) and females (right). Red and blue lines indicate positive and negative correlations with age, respectively. (B) Age trajectories of FC- and EC-based TDA features for the whole brain. The top row shows raw feature values with fitted trajectories; the middle row shows fitted mean trajectories with pointwise 95% confidence intervals; and the bottom row shows age-varying standard deviation trajectories with pointwise 95% confidence intervals. Blue and red curves represent males and females, respectively.

At the higher-order topological scale, TDA-derived features of the whole brain also showed clear but feature-specific age dependence (Figure 7B). Age-related declines were more consistent for EC-based measures, particularly BN-, PE-, and CC-related features, whereas FC-based measures showed more heterogeneous trajectories, including modest declines in PE and CC, a slight increase in ML, and a shallow nonlinear pattern in BN. Male and female trajectories were generally similar in overall shape, but the magnitude of sex separation varied across features and often narrowed with increasing age rather than remaining stably offset throughout adulthood. In several features, the fitted trajectories became progressively closer in later life, and some approached near-overlap or mild crossover. The corresponding trajectories for the cerebrum, cerebellum, diencephalon, and brainstem are shown in Supplementary Figure 6. These four systems showed substantial heterogeneity in the direction, magnitude, and nonlinearity of age-related change. Features that declined at the whole-brain level could show attenuated, flatter, or differently shaped trajectories within individual systems, indicating that whole-brain summaries capture the dominant overall pattern but do not fully represent the anatomical diversity of age-related network organization.

## 4. Discussion

The present study establishes a unified multimodal and multiscale MRI framework for the normative characterization of whole-brain organization in healthy adults, with explicit representation of cerebral, cerebellar, diencephalic, and brainstem regions. By integrating these systems into a shared anatomical space and combining nodal, edge-wise, and higher-order topological analyses, the MIRATS framework addresses a common limitation of conventional cortex-centered neuroimaging. Three main findings emerged. First, deep nuclei occupied distinctive positions within the normative feature space rather than serving as a uniform background to cortical organization. Second, sex-related variation was detectable across nodal, edge-wise, and higher-order topological scales, including within deep systems, but was selective rather than global. Third, age-related variation was widespread yet heterogeneous across feature classes, anatomical systems, and analytical scales.

### 4.1. Deep regulatory systems in the organization of the whole brain

A central aspect of this study is the establishment of a whole-brain normative framework with explicit representation of deep regulatory systems. Human neuroimaging has long been dominated by cortex-centered analytical strategies, whereas small diencephalic and brainstem nuclei have often been underrepresented because of limited spatial resolution, insufficient anatomical contrast, and the lack of standardized templates [12, 49]. With recent advances in high-resolution structural and functional imaging, together with probabilistic mapping of small nuclei, it has become increasingly feasible to measure these regions in vivo and to assess their coupling with cerebellar and cortical systems [8, 50, 51].

Within the normative feature space defined here, diencephalic and brainstem nuclei exhibited systematic profiles that differed from those of cortical and cerebellar regions across morphometric, microstructural, connectional, and topological measures. This observation suggests that deep nuclei occupy distinct positions within whole-brain architecture rather than simply representing small-volume background structures. Such a pattern is biologically plausible, as converging anatomical and physiological evidence points to extensive interconnections among the cerebellum, brainstem, and diencephalon, together with their key roles in arousal, autonomic regulation, respiration, and homeostatic integration [4, 10, 52]. It is also consistent with increasing attention to incomplete cortex-dominant representations of the human brain, particularly in the study of systems that depend critically on lower-level regulatory circuitry [53].

### 4.2. Multiscale representations define a hierarchical normative space

A second major contribution of this study is methodological: it demonstrates that whole-brain organization is inherently hierarchical and cannot be adequately reduced to a single descriptive scale. Regional tissue features quantify local substrate properties, connectivity features characterize pairwise dependencies and system embedding, and topological descriptors capture mesoscale organization that emerges only when the network is considered as a whole. Recent multiscale perspectives argue that these levels should be interpreted as complementary rather than redundant, because each captures partially distinct aspects of neural organization [54, 55].

In the proposed MIRATS framework, nodal measures were most informative for anatomical heterogeneity and age-related tissue variation, whereas FC and EC clarified how regions participate in distributed structural and functional systems. Topological descriptors added a further layer by revealing higher-order organizational patterns that are not fully captured by node-or edge-wise summaries. This interpretation is consistent with recent higher-order connectomic studies showing that topological signatures can improve task decoding, individual identification, and behavioral prediction beyond pairwise connectivity alone [20, 56].

Taken together, these observations suggest that multiscale analysis should not be treated as a set of parallel readouts, but rather as a hierarchical normative language in which regional, connectional, and topological features jointly define the reference space of healthy brain organization. This matters interpretively because a disease effect may manifest as a focal tissue abnormality, an altered interaction pattern, or a deviation that is only apparent at the mesoscale. Normative modeling frameworks provide the conceptual basis for making this distinction, and recent extensions further show how such references can be used to quantify deviation and change over time [57, 58].

### 4.3. Sex-related effects are selective rather than globally uniform

Sex-related effects were detectable across different topological scales, including within diencephalic and brainstem systems, but their magnitude and spatial distribution varied across feature classes. This pattern suggests that sex-related variation in whole-brain organization is selective rather than globally uniform and may depend on the anatomical system and analytical scale under consideration. This interpretation is consistent with evidence that sex-related differences in brain organization are network- and context-specific [59, 60]. Brainstem connectivity differences have been reported to vary not only by sex but also by menopausal status, including reproductive-stage-related variation in connectivity involving the locus coeruleus and periaqueductal gray [61]. More broadly, sex is increasingly understood as a biological variable that interacts with age, hormonal milieu, and chromosomal background, rather than as a nuisance covariate to be removed [62]. The present findings extend this perspective by suggesting that such selectivity also involves deep regulatory systems related to endocrine-autonomic control, arousal, and ascending neuromodulatory signaling. In this sense, the observed effects are better understood as selective modulation of specific systems than as a single global shift across the whole feature space.

### 4.4. Age-related variations are widespread but heterogeneous

Age-related variations were observed across much of the whole-brain feature space, but their expression depended strongly on the type of measure and the anatomical system considered. The nodal features were particularly sensitive to broad age-related tissue differences, consistent with prior literature showing systematic lifespan-related variation in morphometric and microstructural properties [30, 63]. Importantly, the age-related pattern was not confined to the cortex. Evidence indicates that aging engages distributed systems that include the brainstem, cerebellum, and subcortical nuclei in addition to cortical regions [64]. This broader systems perspective is especially relevant for deep structures such as the diencephalon and brainstem, which contribute to sensory relay, sleep–wake regulation, endocrine-autonomic output, and other homeostatic processes. Age-related variation in these systems may therefore reflect not only tissue decline, but also changes in neurochemical regulation, the vascular environment, and neuroendocrine state [65, 66]. Consistent with this view, the present results suggest that different feature classes capture distinct aspects of age-sensitive reorganization, with connectivity- and topology-based measures complementing structural indices of aging. Taken together, these findings support the need for age-related normative references that preserve system specificity and do not reduce deep regulatory circuits to a cortex-centered baseline.

### 4.5. Implications, limitations, and future directions

This work has several implications. Conceptually, this work supports a shift from cortex-centered models toward a whole-brain view in which cerebral, cerebellar, diencephalic, and brainstem systems are all considered essential components of normative brain organization. Methodologically, it demonstrates the feasibility and utility of integrating nodal, edge-wise, and topological measures within a common analytical framework. From a translational perspective, it provides a normative reference that may inform future studies of disorders involving consciousness, respiration, autonomic instability, sleep-wake regulation, and other functions that depend strongly on deep regulatory systems [4, 52].

Several limitations should also be acknowledged. First, although the MIRATS framework explicitly incorporates small deep nuclei, in vivo measurement of these regions remains challenging because of limited spatial resolution, signal heterogeneity, and atlas-related uncertainty [8, 12]. Second, although multimodal and multiscale integration broadens the scope of characterization, it also increases analytical complexity and can complicate interpretation when different feature classes show divergent patterns. Third, the present results characterize normative variation in healthy adults, and further work is needed to determine their generalizability across wider age ranges, multi-site datasets, and clinical populations. Future studies should also test whether disease-related deviations from this normative feature space can improve biological stratification or clinical prediction.

## 5. Conclusion

In conclusion, we developed a unified multimodal and multiscale MRI framework with explicit representation of cerebral, cerebellar, diencephalic, and brainstem regions and applied it to characterize normative whole-brain organization in healthy adults. The findings show that deep nuclei occupy distinctive positions within the normative feature space, and that sex- and age-related variation is selective, anatomically heterogeneous, and expressed across nodal, edge-wise, and higher-order topological scales. By moving beyond conventional cortex-centered representations, this framework provides a more complete basis for describing healthy brain organization and offers a quantitative reference for future studies of disease-related abnormalities involving deep regulatory systems.

