## Supplementary material for "MIRATS framework: Normative multiscale characterization of brain regulatory systems across sex and age using multimodal MRI": Supplementary Materials.docx

This supplementary document provides additional descriptions of the atlas resource, multiscale statistical results, and age-related trajectories referenced in the main text. Supplementary tables report detailed ROI-level, edge-level, and topological statistical results, whereas supplementary figures provide extended visualizations of sex-related and age-related patterns across feature classes and anatomical systems.

**Supplementary Tables**

**Supplementary Table 1.** Regional abbreviations and anatomical descriptions of the 220-region Multiscale Integrated Brain Atlas (MIBA). This table lists all 220 regions of interest included in the MIBA parcellation, together with their abbreviations, full anatomical names, and assignment to one of the four major anatomical divisions: cerebrum, cerebellum, diencephalon, or brainstem. The table serves as the anatomical reference for all ROI-level analyses reported in the main text.

**Supplementary Table 2.** ROI-level sex differences in first-order structural and microstructural features. This table summarizes sex-comparison results for all first-order nodal features, including GMV, GMD, WMV, WMD, FA, MD, AD, and RD, across all 220 ROIs in the midlife subgroup (36--55 years). For each ROI-feature combination, the table reports the test statistic, effect size, and multiple-comparison-corrected significance level. Cohen's d is defined as male minus female, such that positive values indicate higher feature values in males and negative values indicate higher feature values in females.

**Supplementary Table 3.** Edge-wise sex differences in functional connectivity and effective connectivity. This table presents detailed sex-comparison results for second-order edge-wise features, including FC and EC connections that survived false discovery rate correction in the midlife subgroup. For each significant edge, the table reports the source and target regions, t-test statistic, corrected significance value, and effect size direction. These results complement the circular network visualizations shown in Figure 5A.

**Supplementary Table 4.** Sex differences in higher-order topological features across the whole brain and anatomical systems. This table reports sex-comparison results for TDA-derived features computed from FC- and EC-based networks. Reported descriptors include BN, ML, PE, and CC for the whole brain and for each of the four anatomical systems. For each feature, the table includes the group comparison statistic, effect size, and Bonferroni-corrected significance level.

**Supplementary Figures**

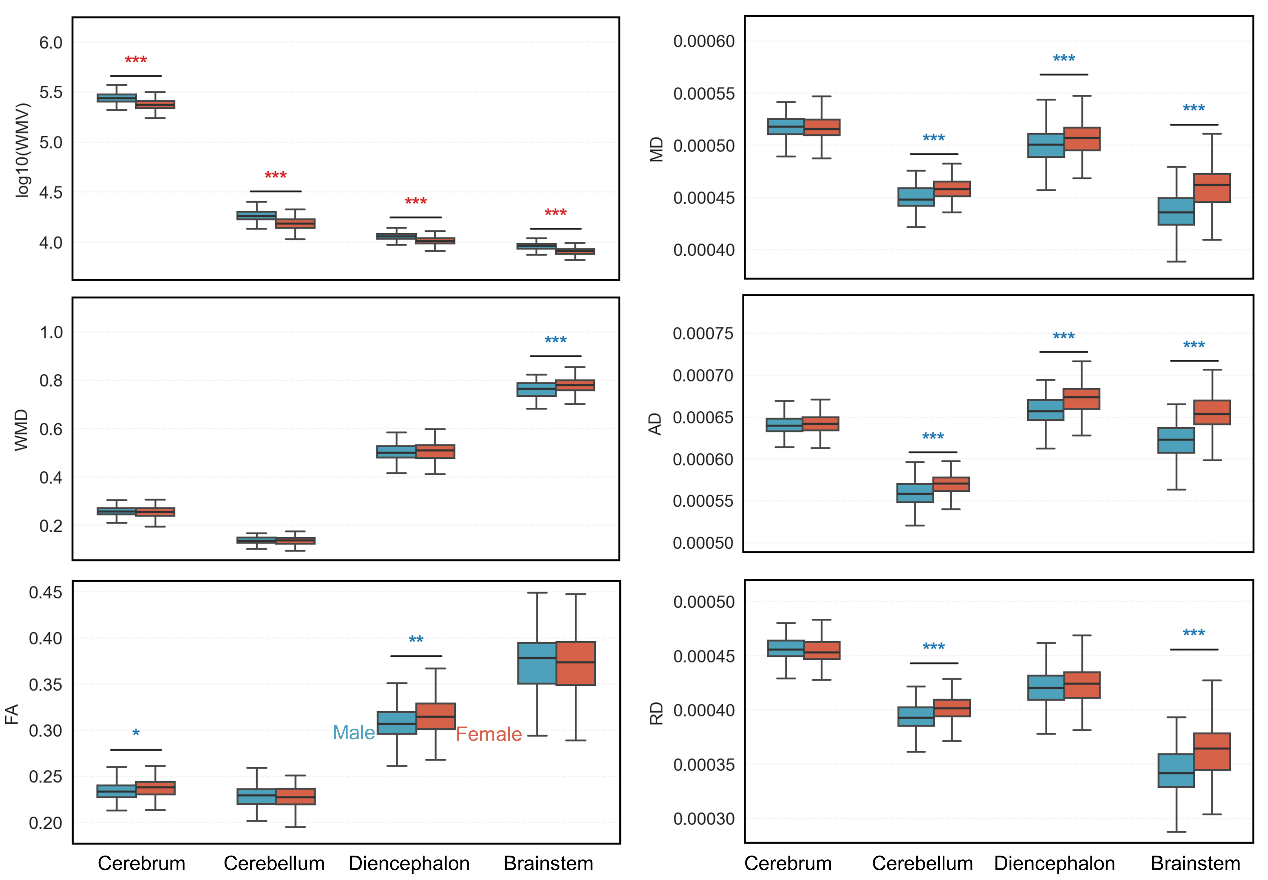

**Supplementary Figure 1.** Boxplots of sex differences in structural and microstructural features not shown in Figure 4. This figure provides ROI-level or system-level boxplot visualizations for the six first-order features not illustrated in the main text figure, namely WMV, WMD, FA, MD, AD, and RD. The plots complement the representative GMV and GMD examples shown in Figure 4B-C and illustrate the direction and magnitude of sex-related variation across anatomical systems. Boxes indicate the interquartile range, center lines indicate the median, and whiskers follow the same convention as in Figure 4.

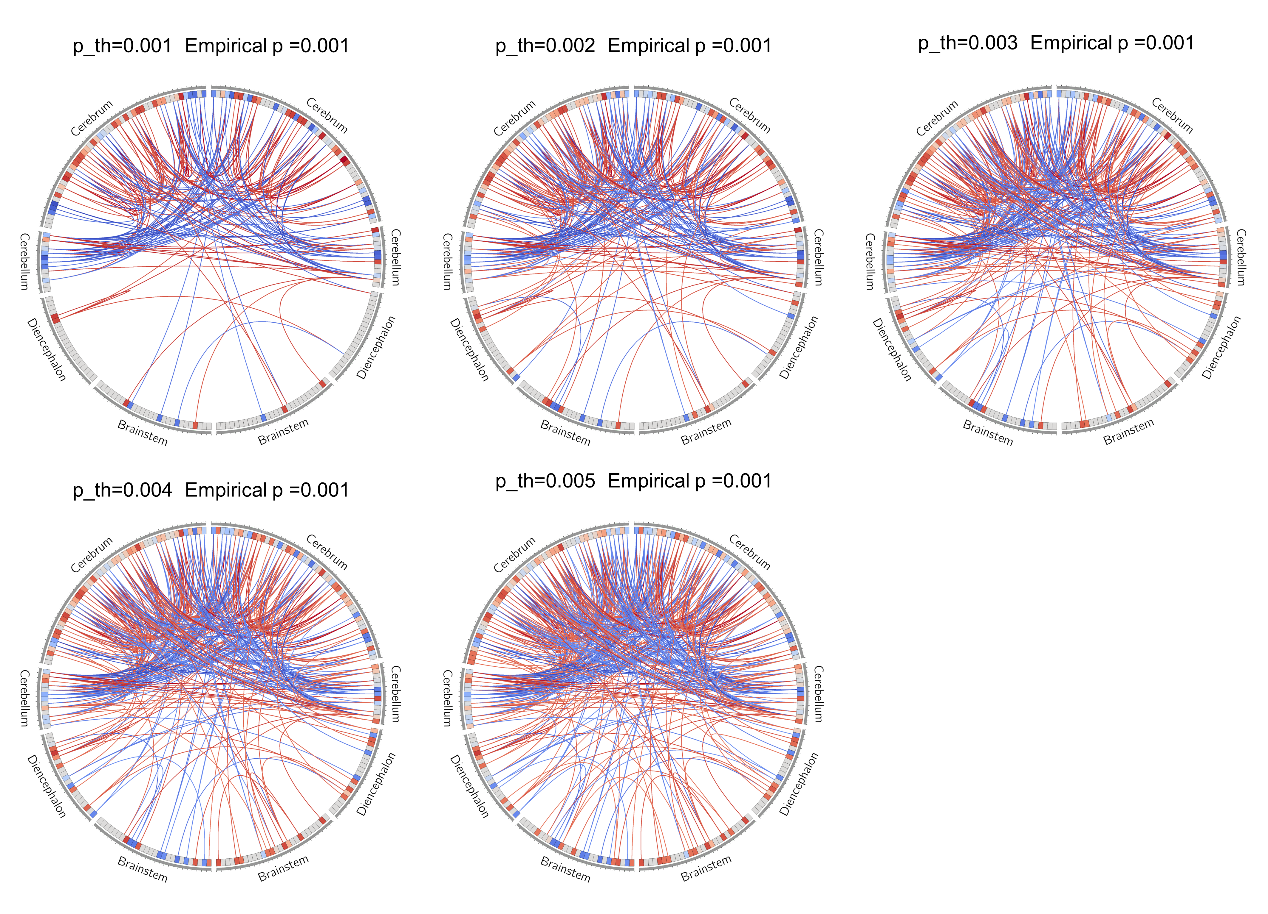

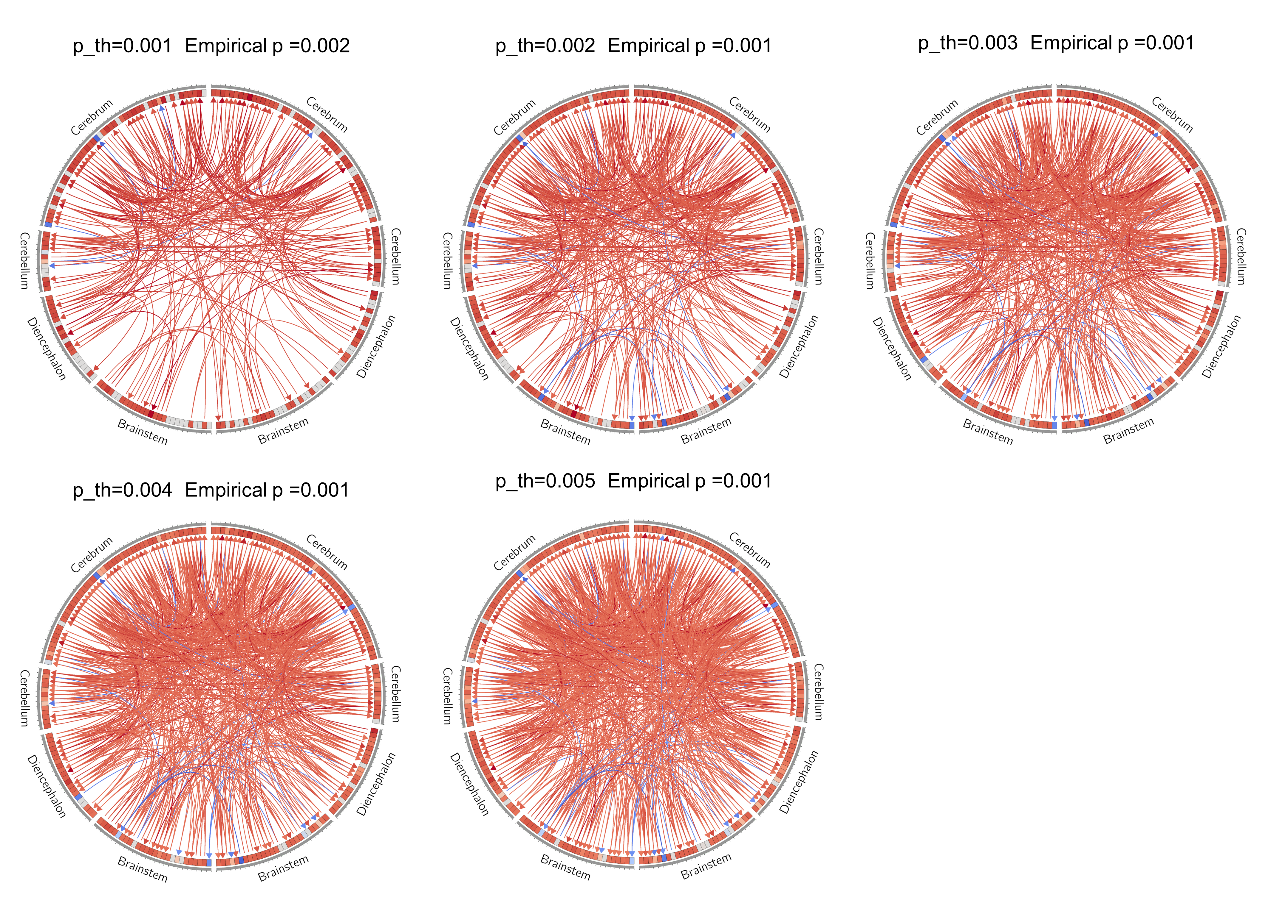

**Supplementary Figure 2.** eNBS-identified connected components showing sex-related differences in FC and EC, respectively. These figures presents the distributed connected components identified by extended network-based statistics (eNBS) for sex-related differences in FC and EC. The figure highlights whether sex effects are expressed as isolated pairwise connections or as coordinated network-level patterns spanning multiple anatomical systems. FC- and EC-based components are shown separately to facilitate comparison of their spatial organization.

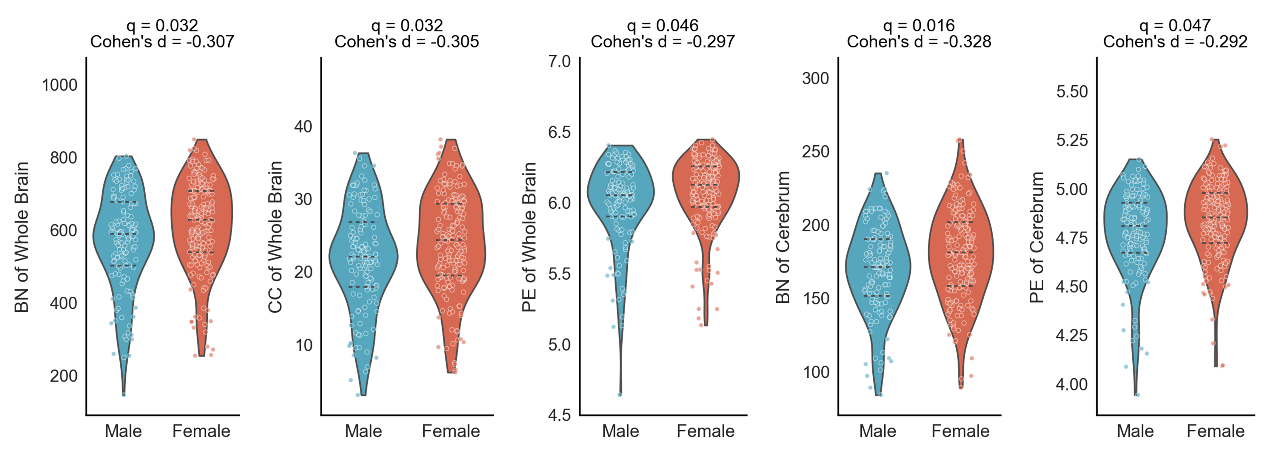

**Supplementary Figure 3.** Violin plots of TDA-derived features showing significant sex differences. This figure shows boxplots for higher-order topological features that exhibited significant sex-related differences after Bonferroni correction. FC-based and EC-based features are displayed separately for the whole brain and the four anatomical systems. These plots visually summarize the direction of group effects and complement the heatmap of Cohen's d values shown in Figure 5B.

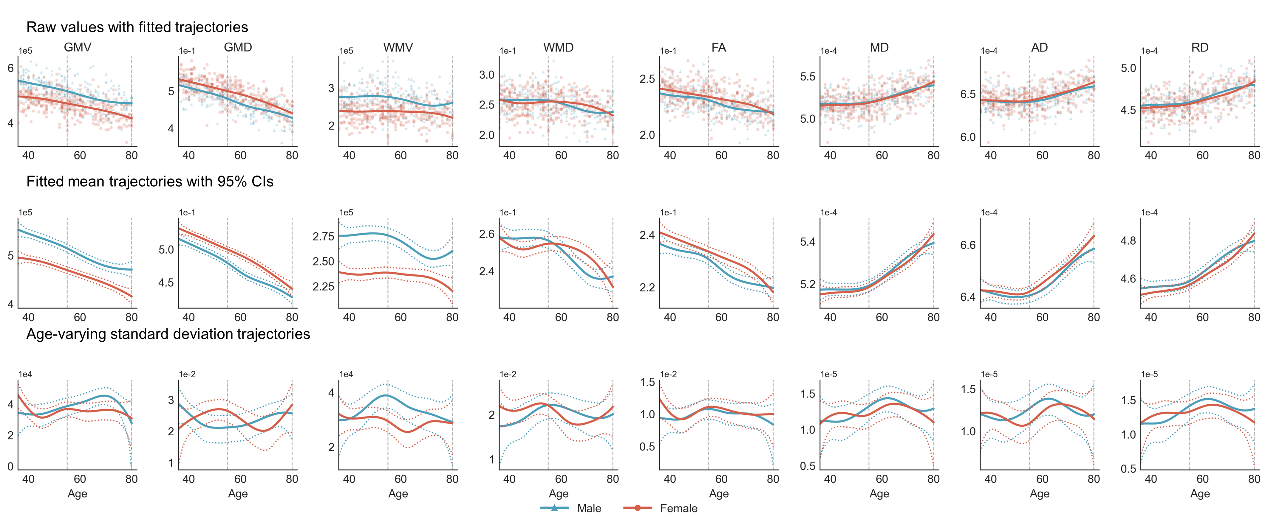

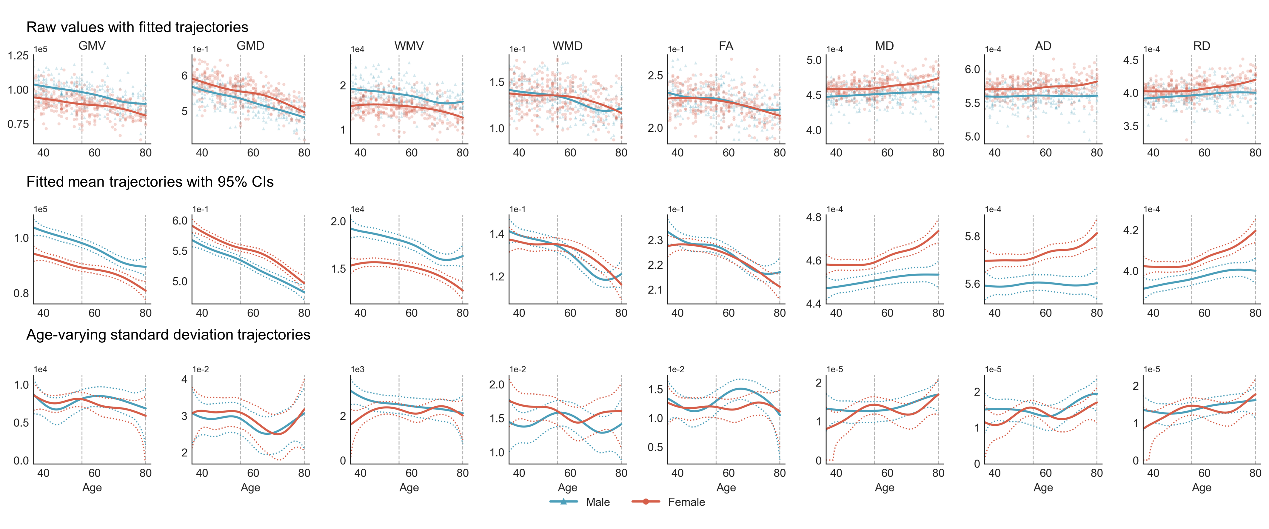

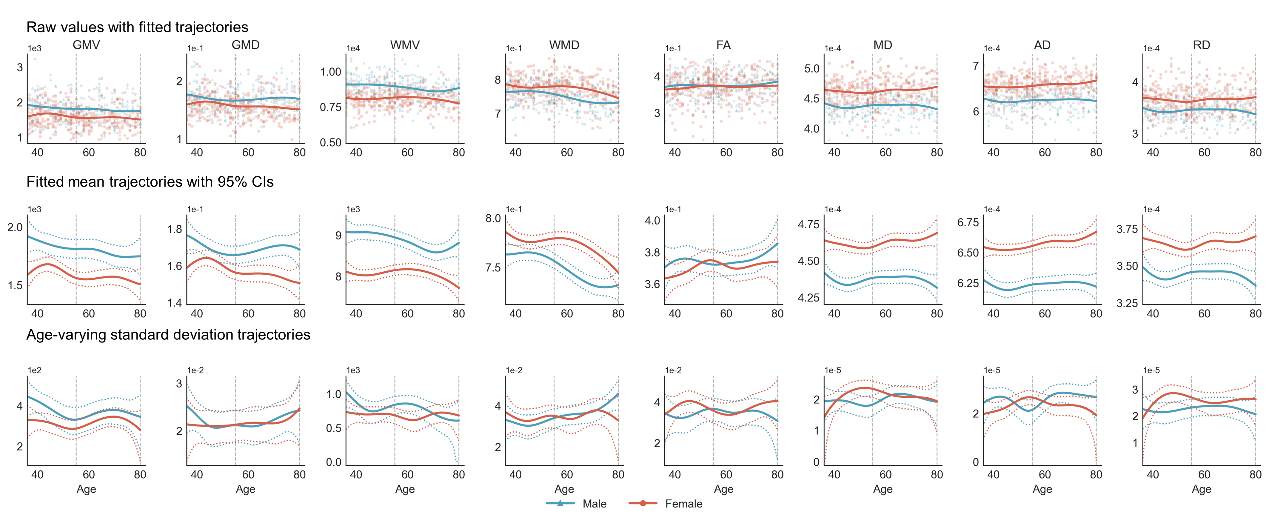

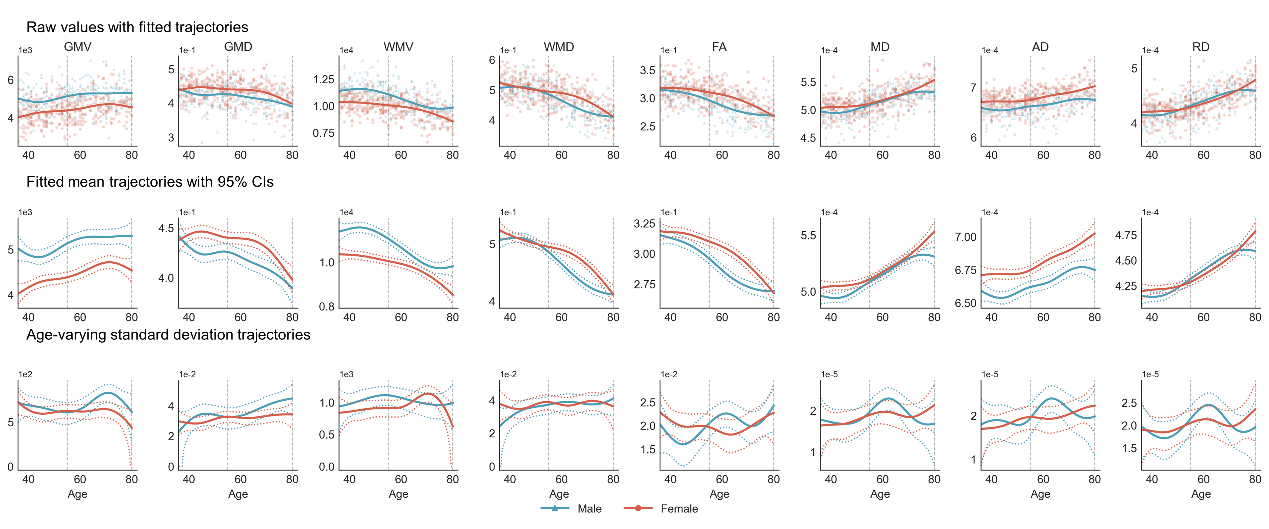

**Supplementary Figure 4.** Age trajectories of structural and microstructural features across the four major anatomical systems. These figures respectively show sex-stratified age trajectories for first-order nodal features within the cerebrum, cerebellum, brainstem, and diencephalon. For each anatomical system, trajectories are shown for GMV, GMD, WMV, WMD, FA, MD, AD, and RD. The top row displays raw values with fitted GAM trajectories, the middle row shows fitted mean curves with pointwise 95% confidence intervals, and the bottom row shows age-varying standard deviation trajectories estimated from a second GAM fitted to squared residuals from the mean model, together with pointwise 95% confidence intervals. Blue and red curves represent males and females, respectively.

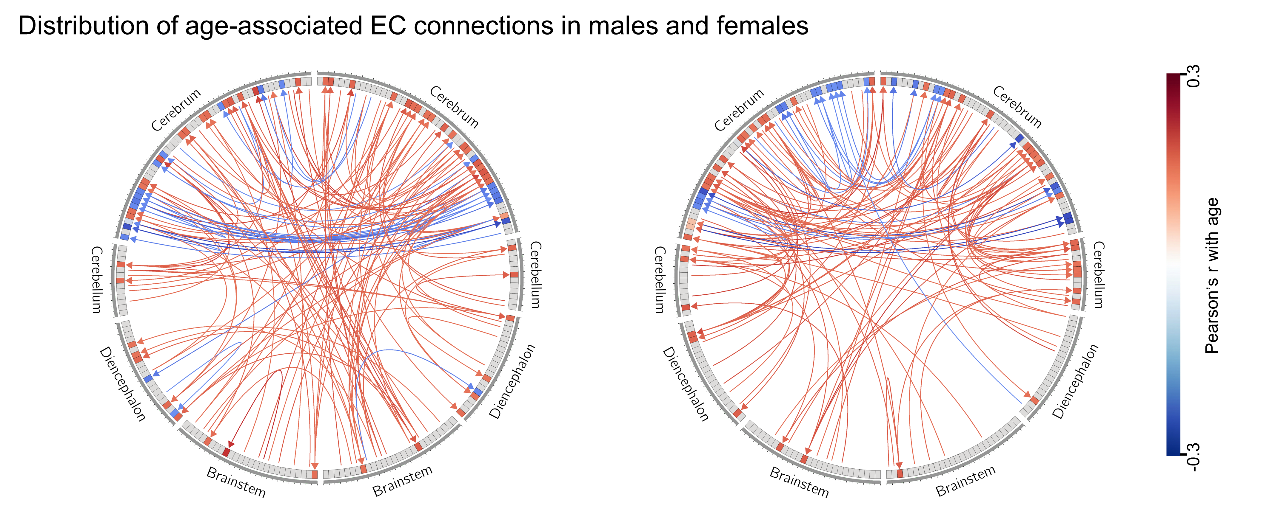

**Supplementary Figure 5.** Age-associated EC patterns in males (left) and females (right). This figure shows the spatial distribution of EC edges displaying age-associated variation in the healthy adult cohort, separately for males and females. As in the FC visualization in Figure 7A, positive and negative associations are displayed using distinct colors. The figure complements the main-text FC results by illustrating whether directed interactions exhibit similar or distinct age-related spatial organization across cortical and deep brain systems.

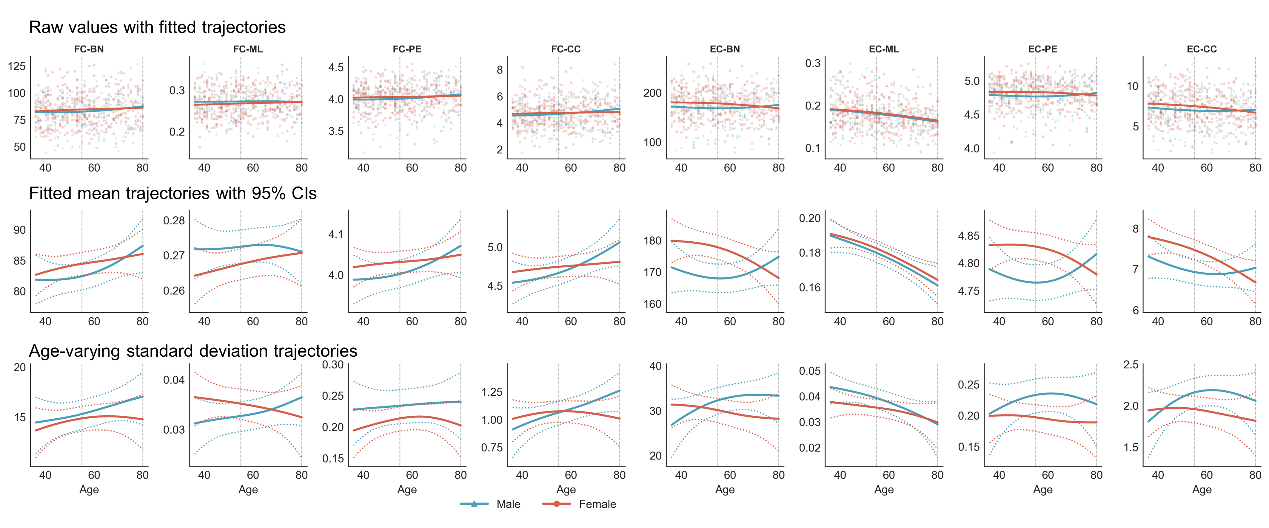

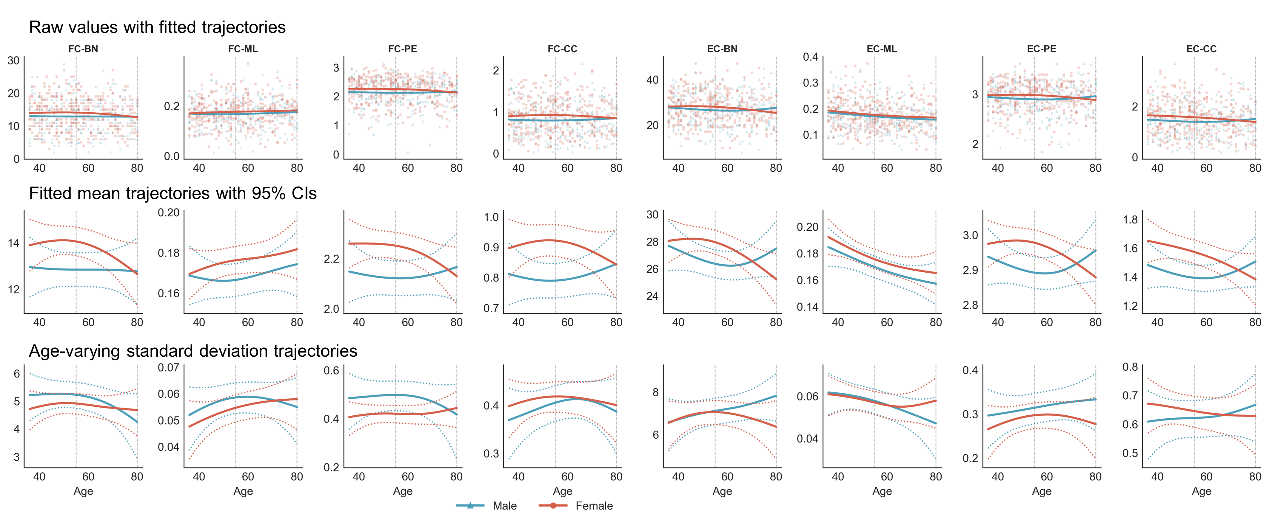

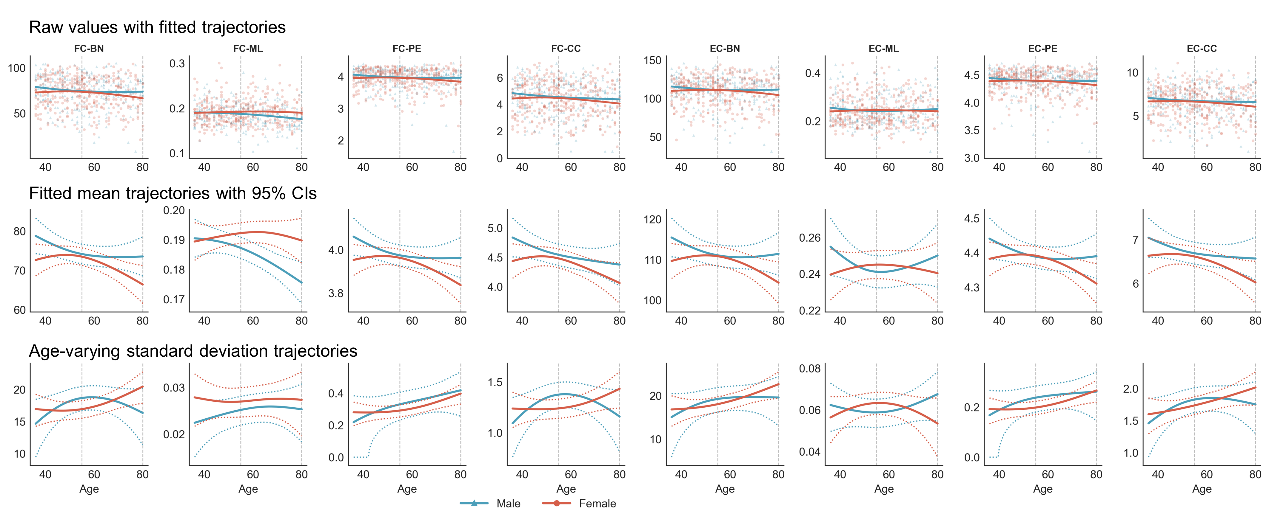

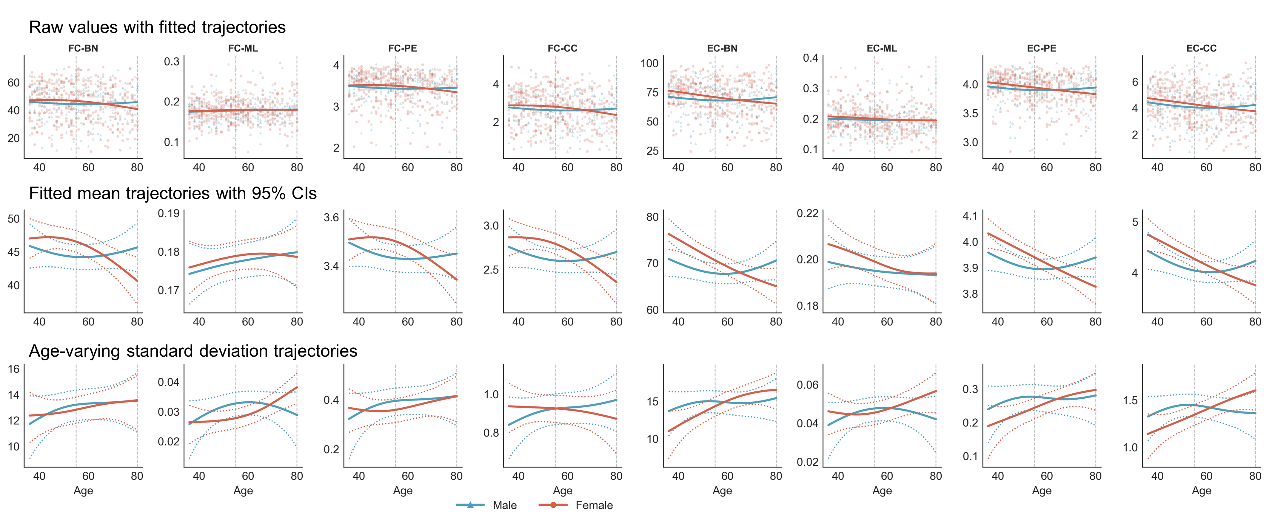

**Supplementary Figure 6.** Age trajectories of TDA-derived features across anatomical systems. These figures respectively present sex-stratified age trajectories of FC-based and EC-based TDA descriptors for the cerebrum, cerebellum, brainstem, and diencephalon. For each system, trajectories are shown for BN, ML, PE, and CC. Raw values with fitted trajectories are shown together with fitted mean curves and pointwise 95% confidence intervals, as well as age-varying standard deviation trajectories estimated from residual-variance smoothing. The figure highlights the heterogeneity of age-related topological change across anatomical systems and complements the whole-brain trajectories shown in Figure 7B.
